# Hybridization In The Late Pleistocene: Merging Morphological and Genetic Evidence

**DOI:** 10.1101/2022.04.20.488874

**Authors:** K. Harvati, R.R. Ackermann

**Affiliations:** Paleoanthropology section, Senckenberg Centre for Human Evolution and Palaeoenvironment, Institute for Archaeological Sciences, Eberhard Karls Universität Tübingen; DFG Centre for Advanced Studies ‘Words, Bones, Genes, Tools’, Eberhard Karls Universität Tübingen; Human Evolution Research Institute, University of Cape Town, South Africa; Department of Archaeology, University of Cape Town, South Africa

## Abstract

This is an exciting time for our understanding of the origin of our species. Previous scientific consensus saw human evolution as defined by adaptive differences (behavioural and/or biological) and the emergence of *Homo sapiens* as the ultimate replacement of non-modern groups by a modern, adaptively more competitive one. However, recent research has shown that the process underlying our origins was considerably more complex. While archaeological and fossil evidence suggests that behavioural complexity may not be confined to the modern human lineage, recent paleogenomic work shows that gene flow between distinct lineages (e.g. Neanderthals, Denisovans, early *H. sapiens*) occurred repeatedly in the Late Pleistocene, likely contributing elements to our genetic make-up that might have been crucial to our success as a diverse, adaptable species. Following these advances, the prevailing human origins model has shifted from one of near-complete replacement to a more nuanced view of partial replacement with considerable reticulation. Here we provide a brief introduction to the current genetic evidence for hybridization among hominins, its prevalence in, and effects on, comparative mammal groups, and especially how it manifests in the skull. We then explore the degree to which cranial variation seen in the fossil record of Late Pleistocene hominins from Western Eurasia corresponds with our current genetic and comparative data. We are especially interested in understanding the degree to which skeletal data can reflect admixture. Our findings indicate some correspondence between these different lines of evidence, flag individual fossils as possibly admixed, and suggest that different cranial regions may preserve hybridisation signals differentially. We urge further studies of the phenotype in order to expand our ability to detect the ways in which migration, interaction and genetic exchange have shaped the human past, beyond what is currently visible with the lens of ancient DNA.

## Current genetic evidence for gene exchange and its effects in hominins

Natural hybridization promotes evolutionary innovation, creating novel and diverse outcomes in subsequent generations, thereby providing a rich substrate on which selection can further act to shape evolutionary trajectories^1, 2^. Since 2010, methodological advances allowing unprecedented, high-resolution insights into ancient genomes have provided increasing evidence for hybridization and resultant gene flow among Late Pleistocene humans. Currently, indications for gene exchange include movement of genes from Neanderthals into early *Homo sapiens* (conventionally called “early modern humans”)^3–8^, resulting in approximately 2-3% Neanderthal ancestry of non-African living modern humans^7^; as well as evidence that *Homo sapiens* contributed to the Neanderthal gene pool as early as 150 to >200ka^9, 10^. Gene flow from Denisovans into the ancestors of modern Asian populations^11, 12^, from Neanderthals into Denisovans^13, 14^, and from some unknown hominin into Denisovans^13^ has also been reported, and the genome of a first generation descendent of a Neanderthal mother and a Denisovan father living ca. 90ka was recently discovered^15^. Finally, genetic exchanges between ancient and recent lineages may have also occurred within Africa^9, 16–21^. Taken together, these studies indicate that gene flow has been multidirectional, was much more common than previously appreciated by most (but see e.g.^22^), and may have been instrumental in structuring genetic diversity across our ancestral lineage over the last half a million years. Given the speed at which new discoveries and methodological breakthroughs are occurring, such as the retrieval of hominin DNA from cave sediments^14^, our expectation is that such evidence will likely continue to accumulate in the future.

Gene flow among hominins has had variable effects, best documented over the last 100K years. These include genetic evidence for some level of introgression affecting phenotypes in a beneficial manner, including those involved in immunity, spermatogenesis, adaptation to low-oxygen contexts, response to ultraviolet radiation, and other traits^23–31^ (but see^32^). For example, Neanderthal genes affecting skin and hair phenotypes are retained in humans living today, (e.g.^27, 31^), suggesting that these genes might have been important in the dispersal and adaptation of people emerging from Africa and migrating into environments inhabited by Neanderthals. In other cases, gene exchange may have been detrimental. For example, the existence of chromosomal regions in living humans devoid of Neanderthal-derived alleles, such as the X-chromosome and genes related to testes and therefore reproduction^27, 31^, suggests that selection may have acted to purge these genes from descendants. Neanderthal alleles present in living people have also been associated with a range of phenotypes considered detrimental in modern (but not necessarily ancient) contexts, including depression, neurodevelopmental disorders, hypercoagulation, altered carbohydrate metabolism, and addiction^27, 29, 33^ (but see^32^). A few recent studies suggest that Neanderthal-derived genetic variation also influences brain phenotypes^29, 34, 35^ and susceptibility to infectious diseases^36, 37^.

Taken together, the genetic evidence to date indicates that gene flow played an important role in shaping the evolutionary fate of our lineage^38^, although its exact effects appear to vary considerably across time, population, and environmental/geographical context.

## Hybridization in extant primates and other mammals and its relevance to hominins

Although the genetic evidence for hybridization in hominins has shifted the prevailing narrative about human origins over the last decade, there was already a growing realisation prior to this (e.g.^39, 40^) that its role may have been underappreciated, based on an increasing understanding of its prevalence across other mammals, including primates. We now know that approximately ten percent of animal species produce hybrids, and occasionally “phylogenetic hotspots” occur in which hybridization rates in animals exceed those seen in plants^41, 42^. Within mammals, hybridization occurs across a wide range of lineages, including (but not limited to) a number of large-bodied terrestrial groups such as bovids^43–45^; bears^46, 47^; cats^48, 49^; canids^50–54^ and primates (see below).

These studies have provided compelling evidence that gene flow impacts the evolutionary trajectory of large-bodied mammals, acting as a particularly strong force for accelerating evolution in novel or changing environmental contexts^1, 2^, a scenario that resonates with the narrative of human origins.

Non-human primates are arguably the most relevant models for human evolution, and there is considerable evidence for hybridization in the wild within all the major lineages at both specific and intraspecific levels including strepsirrhines (e.g.^55, 56^), American monkeys (e.g.^57–61^) and Afro-Eurasian monkeys (e.g.^62–66^). Among these, perhaps the best studied are baboons (genus *Papio)*, which have also repeatedly been put forth as models for human evolution^40, 67^. The six recognised baboon species (or ‘allotaxa’; see^40^) have parapatric ranges, with natural hybridization recorded between the species that are most phylogenetically distant (*Papio ursinus* vs. *P. cynocephalus*), morphologically distinct (*P. ursinus* vs. Kinda baboons), and behaviourally different (*P. hamadryas* vs. *P. anubis*)^68^. Like our own genus *Homo*, *Papio* is the evolutionary product of a radiation that began in non-forested regions of tropical Africa around two million years ago; both genera have inhabited similar regions in Africa and been subject to comparable climatic fluctuations.

Hybridization has also occurred among our closest primate relatives, the apes (Superfamily Hominoidea). It is well-documented among the small-bodied apes (e.g.^69, 70^), and there are also genetic signatures of gene flow both among subspecies (e.g.^71^) and between species^72^ of great apes. One percent of the central chimpanzee genome has been shown to derive from the bonobo^72^, indicating two ancient hybridization events comparable to the admixture seen between modern humans and Neanderthals.

Hybridization can have a wide range of effects on anatomy, behaviour and speciation (see^73^), but it is in its interplay with adaptation that its impact may be most powerful. However, the impact of adaptive introgression can differ even among closely related taxa. For example, in chimpanzees the regions of adaptive introgression are subspecies-specific (e.g. regions involving male reproduction versus immune system)^74^. As a species-specific example, the region around the FOXP2 locus is devoid of introgression in humans^75, 76^, but not in either chimpanzees^77^ or bonobos^78^.

## How does gene exchange manifest itself in skeletal morphology?

Genetic evidence for the effect of gene flow on the hominin skeleton remains limited, despite its importance in linking the genetic and fossil record, as well as potentially understanding the functional implications of skeletal variation. Recent studies suggest that Neanderthal-derived genetic variation influences shape variation in the crania and brains of Europeans living today^34, 35^. In particular, Neanderthal ancestry was found to be associated with a more Neanderthal-like, elongated, cranial and endocranial shape in these Europeans, including morphology in the occipital and parietal regions, as well as differences in brain morphology^34, 35^.

Genetic evidence aside, some researchers have proposed hybrid individuals in the human fossil record on the basis of their morphology. Such proposed hybrids include Lagar Velho 1^79^, Mladeč 5 and 6^80^, Cioclovina 1^81^, Peştera cu Oase 1^82, 83^ and 2^82, 84^, Skhul IV and V^84^, Vindija^85^, Klasies River Mouth^86^, Jebel Irhoud and Mugharet el ‘Aliya in North Africa^86, 87^ and possibly others^88–90^. However, these hypotheses have generally not been possible to test, and the hybrid status of these specimens has been disputed^91–93^ or considered inconclusive^94, 95^. This was mainly due to the lack of clear expectations about hybrid morphology that could be empirically applied to the fossil record (but see^84, 96–99^), but also because of the problem of equifinality, as phenotypes consistent with hybridization, especially “intermediate” morphology, may also be produced through other processes, most importantly by the retention of primitive features. In the face of these shortcomings, admixed status has almost exclusively been recognized on the basis of genetic evidence (as discussed above). However, such evidence is limited in many respects. For example, the application of ancient DNA is constrained due to preservation issues, which can vary from site to site and specimen to specimen, but become particularly severe as we move further back in time or into warmer climates. Additionally, knowledge derived from comparisons among extant genomes can only provide partial insight into the past, given the extinction of many ancient lineages. Therefore, evidence for hybridization present in the skeletal phenotype remains essential to the interpretation of the fossil record, as it can help us to locate such potential events in time and place, and particularly within lineages for which we do not have a genetic record.

The taxon for which most empirical evidence for the effects of hybridization on the skeleton is available is baboons. Studies of baboons have revealed visible perturbations in dental and sutural formation at high frequencies in early generation inter-specific hybrids, as well as atypical expression of some dental traits^1, 96, 97^, suggesting that hybridization breaks down the coordination of early development, although this does not appear to meaningfully affect fitness^84, 97, 100^. These results are consistent with what is seen in the skeletal anatomy of hybrids in other mammalian lineages, including ungulates^43^, rodents^101^, and most recently canids^102^, although they manifest somewhat differently in each taxon. Hybrid baboons also have, on average, crania that are larger than an intermediate value between their parents^96, 97, 103^, with some measurements that are extreme relative to both parents. The production of extreme hybrid phenotypes, or transgressive phenotypes, outside of the range of both parental taxa (in a negative or positive direction) is called transgressive segregation^104^, and in the case of the mammals mentioned above could include atypical traits as well as extreme size/shape.

Large cranial size in hybrids (relative to either a parental midpoint value, or the mean of the largest parent) has also been identified in mice^98, 105–110^ (as well as tamarins^111^). Inter-subspecific mouse hybrids (F1s, F2s, and backcrossed individuals) are typically as large as or larger than the larger parent taxon, with associated size-related shape changes^98, 110^. They also tend to have cranial and mandibular shape variation that is somewhat intermediate to the parents, but more closely resembling the smaller parent (Figure 1), with high levels of heterosis in certain features such as molar length^98, 110^. Later generations (F2, B2) are more variable than first generation hybrids, with backcrosses expectedly moving towards the shape of the parent taxon with which they are hybridizing. These patterns hold for crosses of taxa that hybridize in the wild but have low levels of gene flow and low hybrid fertility; for taxa that hybridize in the wild and produce successful offspring; and for taxa that are geographically separated in nature but nevertheless hybridize under laboratory conditions^98, 110^, making them robust across different scenarios of contact and hybrid fitness. Importantly, the fact that these models show similar patterns for hybridization across both species and subspecies provides a robust model for assessing its impact in taxa where the specific status is debated, such as Neanderthals and *H. sapiens* (See text box “Are Neanderthals Distinct Species and Does It Matter?”).

**Figure 1:**
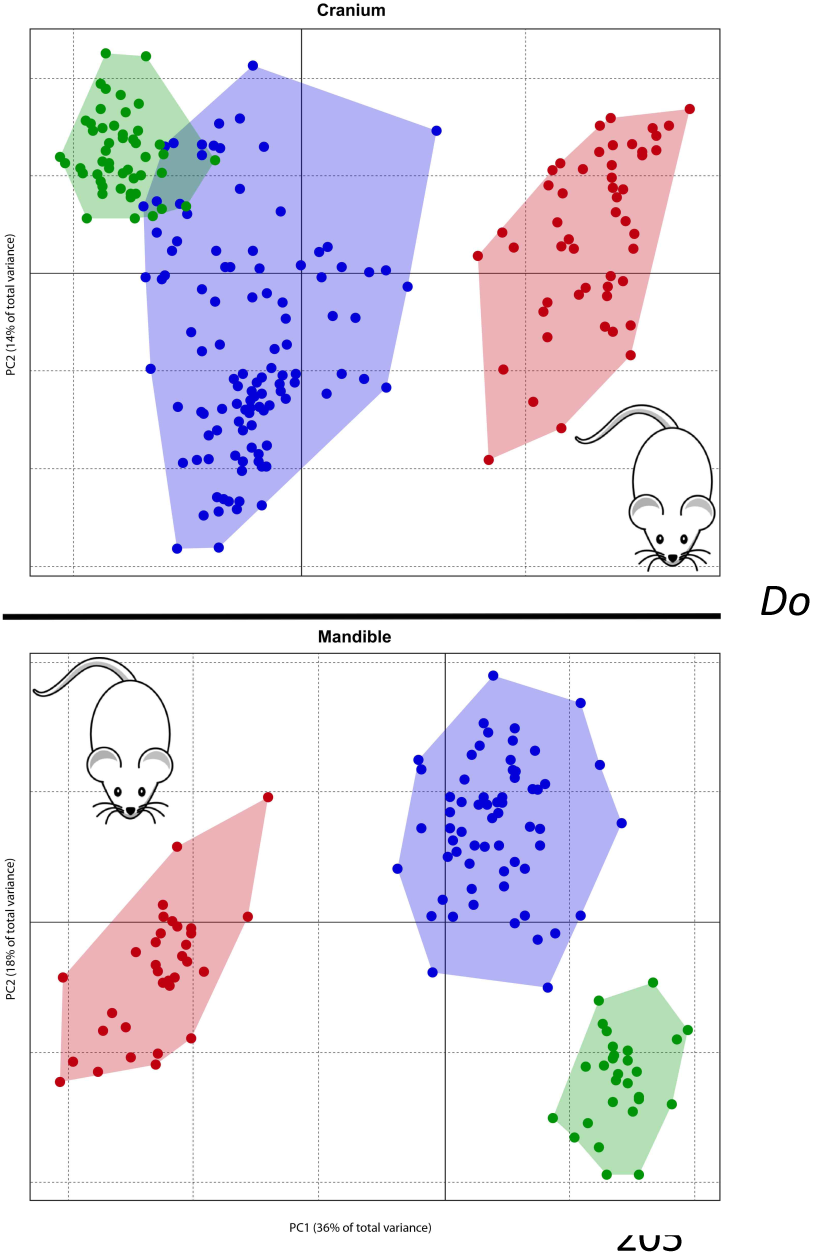
Principal components analysis, mice crania (top) and mandibles (bottom). Red and Green represent parent taxa; Blue represents a pooled sample of first (F1), second (F2), and unidirectionally (with the green parent) backcrossed (B1) inter-subspecific hybrids. Data from^110^.

The studies above have focused on skulls. Unfortunately, considerably less information exists on the effects of hybridization on the postcranial skeleton, outside of the observation from a number of previous studies in mice and primates that hybrids generally exhibit both longer limbs and increased body size relative to parents (e.g.^84, 98, 105, 106, 109, 112–114^), though a new study of macaques suggests that effects of admixture on the pelvis may be relatively small, possibly due to functional or developmental constraints, or relatively minor divergence of the parent taxa (in this case at or possibly below the subspecies level)^115^.

## Do Late Pleistocene Western Eurasian humans fit the morphological predictions of a hybrid sample?

Although current genetic evidence indicates that hybridization occurred repeatedly among Pleistocene hominins, there have been few efforts to link this genetic evidence to morphological evidence from the fossil record itself, despite such a link being key for ascertaining the status and relevance of the bulk of the fossil record (for which genetic data are not available). This is further exacerbated by the fact that our ability to extrapolate from genotype to skeletal phenotype is currently very limited. While it is true that some individuals that show genetic evidence of admixture have limited morphology (e.g.^15^), making establishing these links difficult, this is not the case for other specimens. This lack of discourse between morphology and genetics is detrimental to understanding the dynamics of human evolution in the Late Pleistocene.

Here we explore how insights derived from genetics and model organisms might be applied to the interpretation of the human fossil record. In particular, we examine the patterns of variation in cranial shape across the late Middle to Late Pleistocene, interpreted in conjunction with published genetic and non-metric phenotypic evidence for hybridisation, as discussed further below. The latter evidence consists of sutural and dental developmental anomalies comparable to what has been observed in comparative studies on hybridization and its effects on the phenotype. Even though admixture between hominin lineages has been demonstrated outside of Western Eurasian contexts, we focus specifically on Neanderthals and early *Homo sapiens*, i.e. hominins from Western Eurasia and Africa, in order to limit the scope of this inquiry. In keeping with current consensus, we consider these taxa to represent distinct lineages evolving in large part independently (see, e.g.^116^). However, in recognizing the important limitations of species concepts and their application to the fossil record, as well as long standing disagreements on Neanderthal alpha taxonomy, we avoid the term *Homo neanderthalensis*, while using *Homo sapiens* to refer to extant humans and their ancestors in the late Middle and Late Pleistocene, following recent literature (e.g.^116–118^). We consider the following questions: Do Late Pleistocene Eurasian *H. sapiens* as a sample match the anatomical expectations, based on the mammal comparative data, for a Neanderthal-early *H. sapiens* hybrid population spanning multiple generations? Among individuals for whom genetic data are available, does a higher level of Neanderthal ancestry co-occur with Neanderthal-like morphology or with developmental abnormalities? How does hybrid status manifest itself in different aspects of cranial shape and size, and are these skeletal indicators useful predictors of admixture in samples where no genetic evidence is available?

Our sample comprises late Middle and Late Pleistocene (roughly corresponding to MIS 7-2) fossil human specimens from Europe, Africa and the Middle East assigned to Neanderthals and *H. sapiens* (Table 1; Figure 2), including, but not limited to, individuals that are genetically known and morphologically proposed hybrids. We chose an upper age limit of MIS 7 because the suites of diagnostic morphological features of both Neanderthals and modern humans were largely established by this time (see e.g.^116, 119^). In order to frame our study in a manner that is consistent with studies from model organisms (e.g. see Figure 2), the Neanderthal portion of this sample is considered as representative of one of the “parental” (unhybridized) populations. We cannot rule out *H. sapiens* ancestry in individual Neanderthals, though to date evidence for introgression of *H. sapiens* genes into Neanderthals is more limited than the reverse. For the second parental population, because of the poor representation of penecontemporaneous African early *H. sapiens* in our data set and in the fossil record generally, we consider a pooled sample of ancient and recent sub-Saharan Africans, expected to have no or minimal Neanderthal ancestry (see^9^), as a proxy for early *H. sapiens* anatomy. A few individuals from this time period whose attribution to early *H. sapiens* is uncertain (i.e. Omo 2, Eliye Springs) have also been included in some of our analyses, but they are labelled and discussed separately. The recent sub-Saharan African portion of the pooled *H. sapiens* sample is represented by three datasets of individuals from eastern and southern Africa (face: N=15; hemimandible: N=14; posterior cranial profile: N=15). We will refer to this sample as African *H. sapiens*. We recognise that the inclusion of this small sample of extant sub-Saharan Africans is not ideal given the potential effects of recent and ancient demographic processes, as well as the possibility of admixture in deeper time (e.g.^9, 16–21^). We hope to mitigate such effects to the extent possible by combining the few available ancient individuals with our recent African sample and limiting the extant sample to sub-Saharan Africa, thereby reducing the likelihood of admixture from Neanderthals. We cannot rule out the possibility that introgression from other non-Neanderthal “ghost lineages” might be present in the African sample, or the late survival of archaic lineages not directly ancestral to *H. sapiens* in our ancient African samples. Specimens explicitly proposed as such possible hybrids (e.g. the Iwo Eleru calvaria) have been excluded from our analyses. The results presented below, which largely position the ancient African samples within the same shape space as the modern ones (and distinct from Neanderthals), support these choices. All other fossils are considered together as Pleistocene (non-Neanderthal) western Eurasians, which we will refer to as Eurasian *H. sapiens*, and as such are potentially admixed, and the primary focus of this study. The Apidima 1 specimen from Greece, whose attribution to early *H. sapiens* is uncertain^117^, is included as a possible Eurasian early *H. sapiens* but labelled separately.

**Figure 2:**
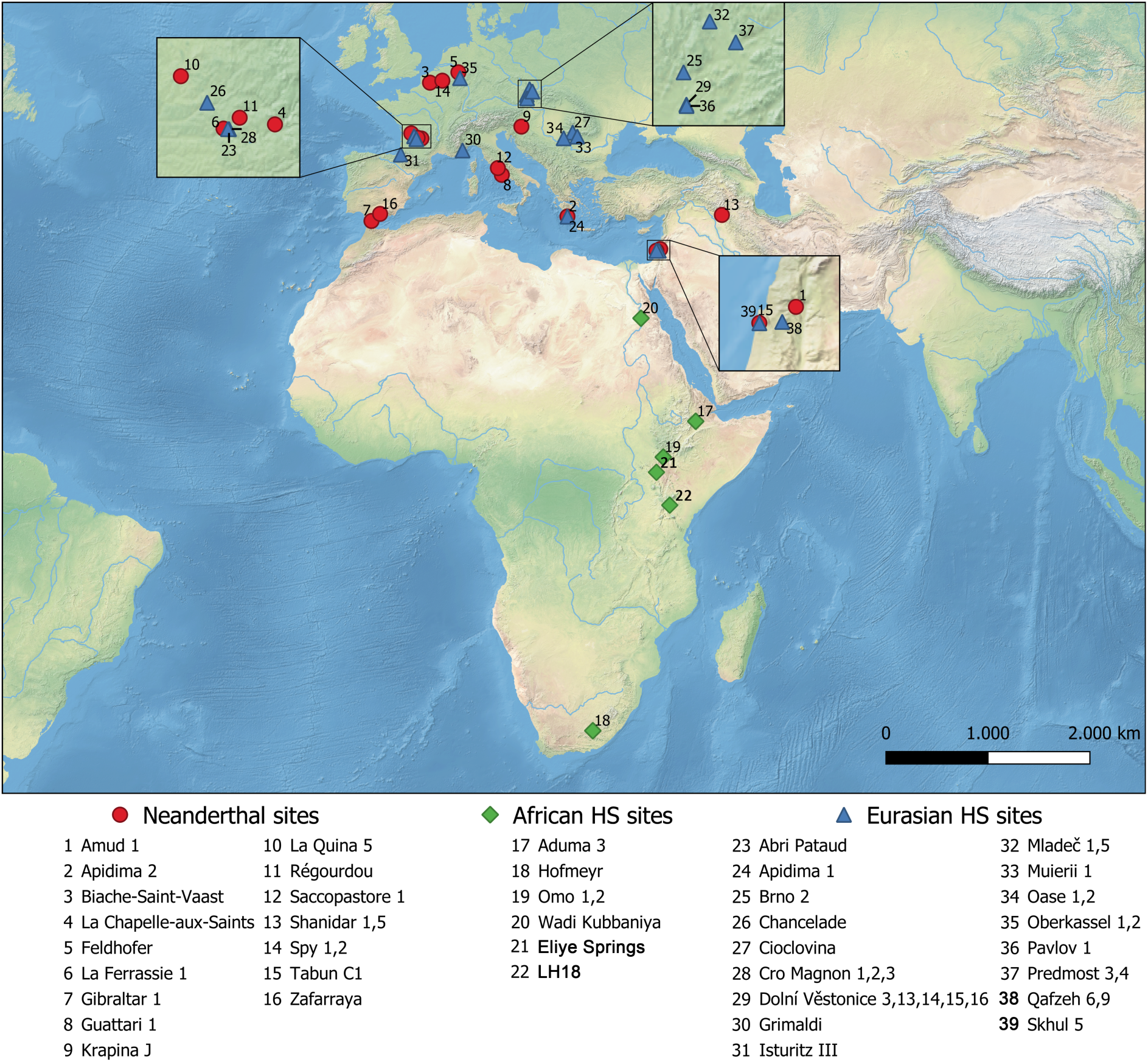
Localities for Pleistocene fossil hominin specimens used in the analyses.

**Table 1:** Hominin samples used in analyses. Dates are reported in calendar years unless otherwise stated

|  | Face | Mandible | Neurocranium | Apx geological age (ka) | Reference |
| --- | --- | --- | --- | --- | --- |
| **Neanderthal** |  |  |  |  |  |
| Amud 1 |  | x | x | 55-60 | 153 |
| Apidima 2 | x |  |  | ca. 170 | 117 |
| Biache St Vaast |  |  | x | ca. 250 | 154 |
| La Chapelle-aux-Saints | x |  | x | 47 / 56 | 155 |
| Feldhofer |  |  | x | 40 | 156 |
| Ferrassie 1 | x | x | x | 43-45 | 157 |
| Gibraltar 1 | x |  |  | ca. 50 | 158 |
| Guattari 1 | x |  | x | 50-60 | 159,160 |
| Krapina J |  | x |  | 140-120 | 161 |
| La Quina 5 |  | x | x | >48 (OIS 3-4) | 162 |
| Regourdou |  | x |  | MIS 4-5 | 163 |
| Saccopastore 1 |  |  | x | 295-220 | 164 |
| Shanidar 1 | x | x |  | 46-50 Uncalibrated | 165 |
| Shanidar 5 | x |  |  | 46-50 Uncalibrated | 165 |
| Spy 1 |  |  | x | 40.6-44.2 cal BP | 166 |
| Spy 2 |  |  | x | 40.6-44.2 cal BP | 166 |
| Tabun C1 |  | x |  | 130-100 | 167,168 |
| Zafarraya |  | x |  | ca 30-46, >46 | 169,170 |
| ***Homo sapiens* Late Middle – Late Pleistocene Africa** |  |  |  |  |  |
| Aduma 3 |  |  | x | 79-105 | 171 |
| Hofmeyr | x |  |  | 36 | 172 |
| Eliye Springs |  |  | x | Middle/Late Pleistocene | 173,174 |
| LH 18 |  |  | x | 120+/-30 | 175,176 |
| Omo 1 |  |  | x | 233+/-22 | 177,178 |
| Omo 2 |  |  | x | 195+/-5 | 178 |
| Wadi Kubbaniya | x | x |  | ca. 20 | 179 |
| ***Homo sapiens* Late Middle – Late Pleistocene Eurasia** |  |  |  |  |  |
| Abri Pataud 1 | x | x | x | 28-26 (22 uncalibrated) | 180 |
| Apidima 1 |  |  | x | ca. 210 | 117 |
| Brno 2 |  |  | x | 23.7 uncal (ca. 28.5 cal BP) | 181 |
| Chancelade | x |  | x | 18 | 182 |
| Cioclovina |  |  | x | ca. 33 | 183 |
| Cro Magnon 1 | x |  | x | ca 30 | 184 |
| Cro Magnon 2 | x |  | x | ca 30 | 184 |
| Cro Magnon 3 |  |  | x | ca 30 | 184 |
| Dolní Vestonice 13 | x | x | x | ca. 31 | 185 |
| Dolní Vestonice 14 | x | x |  | ca. 31 | 185 |
| Dolní Vestonice 15 | x | x | x | ca. 31 | 185 |
| Dolní Vestonice 16 | x | x | x | ca. 30 | 185 |
| Dolní Vestonice 3 | x | x | x | undated | 185 |
| Grimaldi | x | x |  | 25 uncal (ca. 29.5 cal BP) | 186 |
| Isturitz III |  | x |  | Upper Paleolithic | 187 |
| Mladec 1 | x |  | x | 35-36.5 | 188 |
| Mladec 5 |  |  | x | 35-36.5 | 188 |
| Muierii 1 | x | x | x | ca. 35 | 81 |
| Oase 1 |  | x |  | ca. 40.5 | 189 |
| Oase 2 | x |  | x | ca. 40.5 | 82 |
| Oberkassel 1 |  | x |  | 12 uncalb (ca. 14.2 cal BP) | 190 |
| Oberkassel 2 |  | x |  | 12 uncalb (ca. 14.2 cal BP) | 190 |
| Pavlov 1 |  |  | x | ca. 30 | 185 |
| Predmost 3 | x |  | x | 27-29 | 191 |
| Predmost 4 | x |  | x | 27-29 | 191 |
| Qafzeh6 | x |  | x | 100-130 | 168 |
| Qafzeh9 | x | x | x | 100-130 | 168 |
| Skhul 5 |  | x | x | 100-130 | 168 |

We expect that the morphological datasets investigated will differentiate between Neanderthals and African *H. sapiens*, reflecting their commonly accepted status as distinct lineages. Further, we make a series of predictions aimed at determining whether the Eurasian *H. sapiens* sample shows patterns of variation consistent with hybridization between African *H. sapiens* and Neanderthals. As discussed briefly above, empirical research on hybridization in primates, mice, and a handful of other mammals predicts that an admixed sample should contain: (1) individuals with “mixed” or intermediate morphologies somewhere between their parental (un-admixed) taxa, (2) individuals with developmentally atypical traits not seen (or seen at extremely low frequency) in the parental taxa (especially dental or sutural anomalies), and/or (3) individuals that are transgressive in shape or size relative to the parental taxa^1^; these characteristics will frame our morphological expectations. Taken together, they are expected to result in hybrid populations that are more diverse than parental groups. The consistency of these findings across taxa and generations within taxa^1, 97^ support the use of this general pattern for determining hybrid status in the fossil record.

In assessing the question of hybridisation between early *H. sapiens* and Neanderthals, some additional dynamics also need to be taken into account. Introgression from Neanderthals into *H. sapiens* occurred at a low level, likely mediated by differences in population size (with *H. sapiens* considerably larger; e.g.^8, 120^) as well as directionality of backcrossing and possible reduced hybrid fitness (e.g.^121–123^). Moreover, any sample is certain to be comprised of multi-generational recombinants, rather than first generation hybrids. As a result, not all specimens in our Eurasian *H. sapiens* sample are expected to be admixed, and those that are will likely be represent individuals with substantially more African than Neanderthal ancestry components. Indeed, all Upper Paleolithic Eurasian specimens for which genetic information is available show evidence of Neanderthal admixture at least as great as that observed in modern non-Africans, but specimens with recent Neanderthal ancestry are rare^6, 124^. This differs from the studies of model organisms which focus primarily on early generation hybrids. Furthermore, our analyses focus on specific aspects of skull anatomy (mandibular, facial, neurocranial) in order to maximize samples (see below), and therefore our datasets do not replicate exactly the model organism studies that generally examine size/shape of overall cranial morphology (in addition to key non-metric traits). As a result of these factors, we expect to find substantial overlap between the African *H. sapiens* ’parental’ population and a Eurasian admixed sample, with some individuals plotting as expected for hybrids, i.e. intermediate, atypical, or transgressive. Alternatively, if no admixture occurred, or if such admixture does not manifest on the aspects of cranial morphology investigated here, the Eurasian *H. sapiens* sample would be expected to largely conform to the patterns shown by the African *H. sapiens* ‘parental’ population. However, it must be stressed again that an important complicating factor in these assessments is the problem of equifinality, i.e. that similar morphologies can result from different processes. Some of the predictions outlined above for admixture may also apply to other evolutionary processes, such as, for example, the retention of primitive features, or selection for specific phenotypes under particular environmental conditions leading to convergence. The results presented here, therefore, must be interpreted with caution.

Due to the fragmentary nature of the fossil record, individuals are generally not fully preserved, and different individuals are often represented by different parts of the skeleton. In order to include as many fossils as possible, we evaluated three anatomical regions: the face, the hemimandible, and the posterior cranial profile (midsagittal profile). Our data were all collected previously by one of us (KH), and consist of three-dimensional landmarks and semilandmarks, processed with Procrustes superimposition and semilandmark sliding (in the case of the posterior cranial profile), and analysed using Principal Components Analysis (PCA). The datasets were specifically designed to capture salient morphological features that are widely considered Neanderthal or *H. sapiens* derived traits in the respective anatomical regions and are routinely used for taxonomic identification (see e.g.^125–127^). However, they may be affected differentially by different evolutionary processes. For example, facial and mandibular traits may be influenced by selection resulting from environmental factors, such as climate or diet (e.g.^128–130^), with facial morphology also possibly affected by stabilizing selection due to its importance in species recognition^131^. In contrast, neurocranial shape is proposed to track neutral evolutionary changes and population history more closely^128^, and has been linked to Neanderthal genetic ancestry in modern Europeans^35^. Our mandibular and facial datasets, therefore, may be expected to reflect a hybridization signal less clearly than our midsagittal profile dataset. Data indicating the presence of non-metric skeletal abnormalities, and genetic information on % Neanderthal ancestry, where available, were compiled from the literature and integrated in our figures and discussion. For each dataset, we also developed a shape index by calculating an axis between the mean Neanderthal and mean African *H. sapiens* shapes and projecting all Eurasian *H. sapiens* onto it^35, 117^. Results are presented in Figures 3-5.

**Figure 3:**
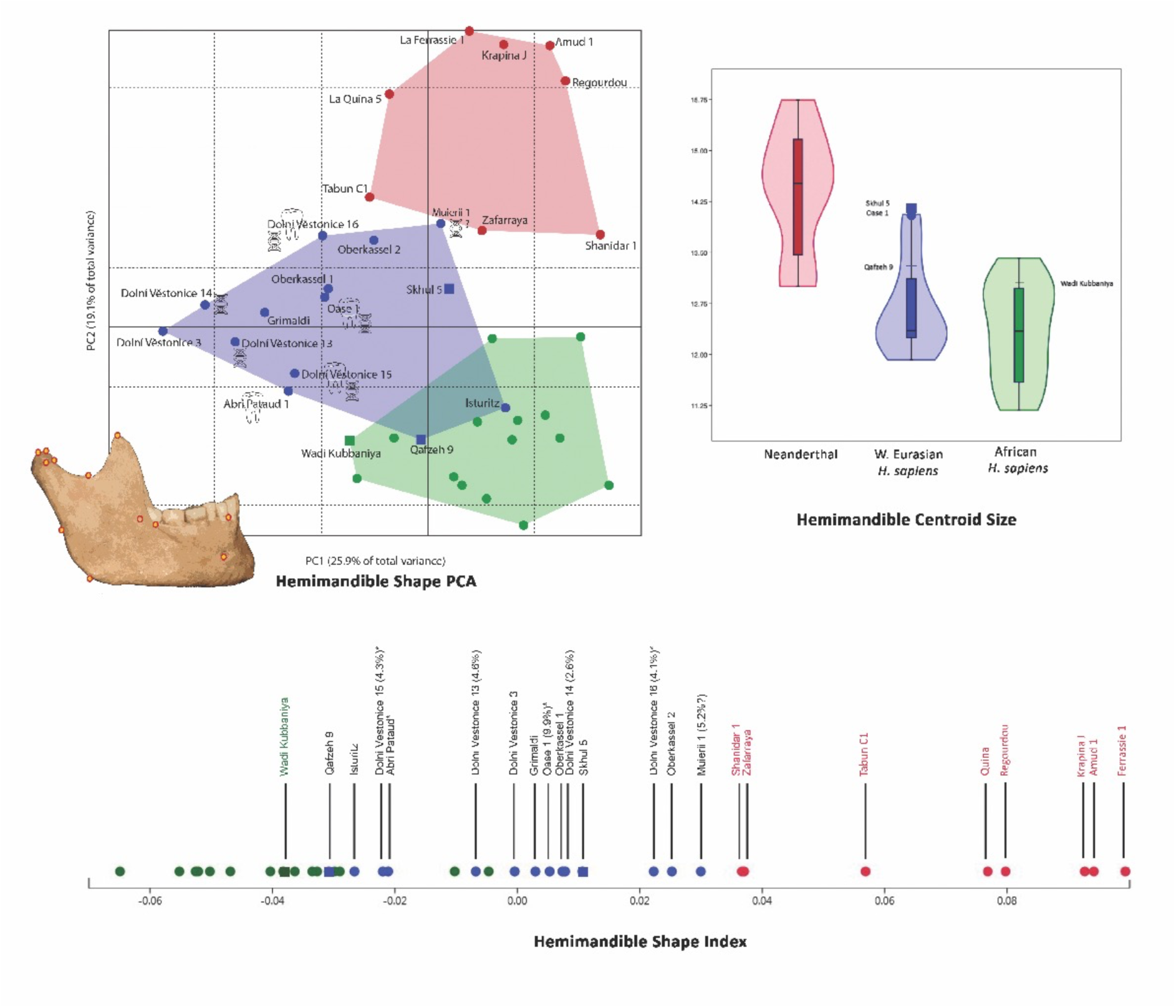
Hemimandible analysis. Top left: Principal Components Analysis, landmark dataset shown on modern human mandible model. Top right: Violin plot, centroid size by group, minimum to maximum values, superimposed box plot showing median and 25-75 % quartiles. Bottom: Shape index. Red: Neanderthals; Green: African recent *H. sapiens* (dots), Pleistocene specimen Wadi Kubbaniya (square); Blue: Eurasian Late Pleistocene *H. sapiens* (dots), early *H. sapiens* (square).

Individuals with genetic evidence for hybridisation are marked with a DNA symbol in the PCA plot, with % Neanderthal ancestry given in the shape index. Individuals with atypical dental or sutural variation as reviewed in^132^ are marked with a TOOTH symbol in the PCA plot and with an (*) in the shape index.

In the hemimandible analysis, the Eurasian *H. sapiens* sample broadly conforms to our expectations for a hybridised sample, occupying shape space somewhat intermediate to the Neanderthals and the African *H. sapiens* in the PCA, with transgressive individuals (Figure 3). This is a similar pattern to that observed in mouse hybrids relative to parental lineages. Although intermediate, the Eurasian *H. sapiens* sample is largely outside of the range of variation seen in either Neanderthals or African *H. sapiens*, overlapping only minimally with African *H. sapiens* in the PCA. Moreover, all the individuals with genetic or morphological signatures of hybridization sit outside of the range of variation in the “parental” samples (neither African *H. sapiens*, nor Neanderthal), with many of them transgressive on PC1. The Eurasian *H. sapiens* sample is also partly intermediate, though closer to the African *H. sapiens*, in centroid size and in the shape index.

However, neither the percentage of Neanderthal genetic ancestry, where known, nor the incidence of developmental abnormalities, appear to follow a clear relationship with Neanderthal-like morphology or with the shape index values. A case in point is the Oase 1 mandible. This individual is known to have approximately 10% Neanderthal ancestry - equivalent to a Neanderthal ancestor four to six generations previously^4, 133^ – and is currently the earliest generation Neanderthal-modern human hybrid known. Oase 1 shows very large overall size (one of the two largest *H. sapiens* mandibles in centroid size in our sample) and megadont lower third molars^83, 132^, consistent with its hybrid status. Yet its mandibular shape index value is less Neanderthal-like than other specimens with known smaller Neanderthal genetic components (Figure 3). Indeed, Muerii 1 (although there is no genome data available on Muierii 1, it may represent the same individual as Muierii 2 with 5.2% Neanderthal ancestry^4^), Oberkassel 2 and Dolní Vestoniče 16 fall closest to Neanderthals in the mandibular shape index.

A similar PCA pattern is shown by the posterior cranial profile analysis (Figure 4), although the separation between the African *H. sapiens* and Neanderthal convex hulls is smaller, a phenomenon largely driven by the position of Omo 1. Although the Eurasian *H. sapiens* sample is again intermediate between Neanderthals and African *H. sapiens*, it shows much more overlap with both ‘parental’ ranges, and especially with the African sample, indicating that a large proportion of these Eurasian specimens display *H. sapiens*-like shape, while some are more Neanderthal-like (and some intermediate / transgressive). This dataset essentially investigates a single – albeit very important – feature, the outline of the posterior part of the cranium in lateral view. A rounded cranium is considered a derived feature for modern humans, and recent work has linked a relatively reduced globularity of the parietal and occipital bones in modern Europeans to Neanderthal genetic ancestry and even to the presence of specific Neanderthal alleles^35^. Our shape index of the posterior cranial profile, encompassing the midsagittal outline of the parietal region and the upper occipital, might reasonably be considered as a proxy for an important aspect of the ‘globularisation’ index calculated by^35^. The overall observed pattern of separation between our Neanderthal and African *H. sapiens* samples is consistent with that described by^35^, with the exception of Omo 2 – and, to a lesser extent, Omo 1. These specimens differ from all other African ones in that they plot within the Neanderthal convex hull (Omo 2) or relatively close to it (Omo 1). Their position on the plot may indicate high levels of variation and population structure in early *H. sapiens*, as has been argued previously (e.g.^134, 135^). Alternatively, Omo 2, the only African specimen overlapping with Neanderthals and showing a Neanderthal-like shape index, may not represent an early *H. sapiens* (see e.g.^116^). The remaining early African *H. sapiens* or possible *H. sapiens,* including LH18, Eliye Springs and Aduma 3, plot with the African sample, with all but Aduma 3 overlapping with the Eurasian *H. sapiens* range.

**Figure 4:**
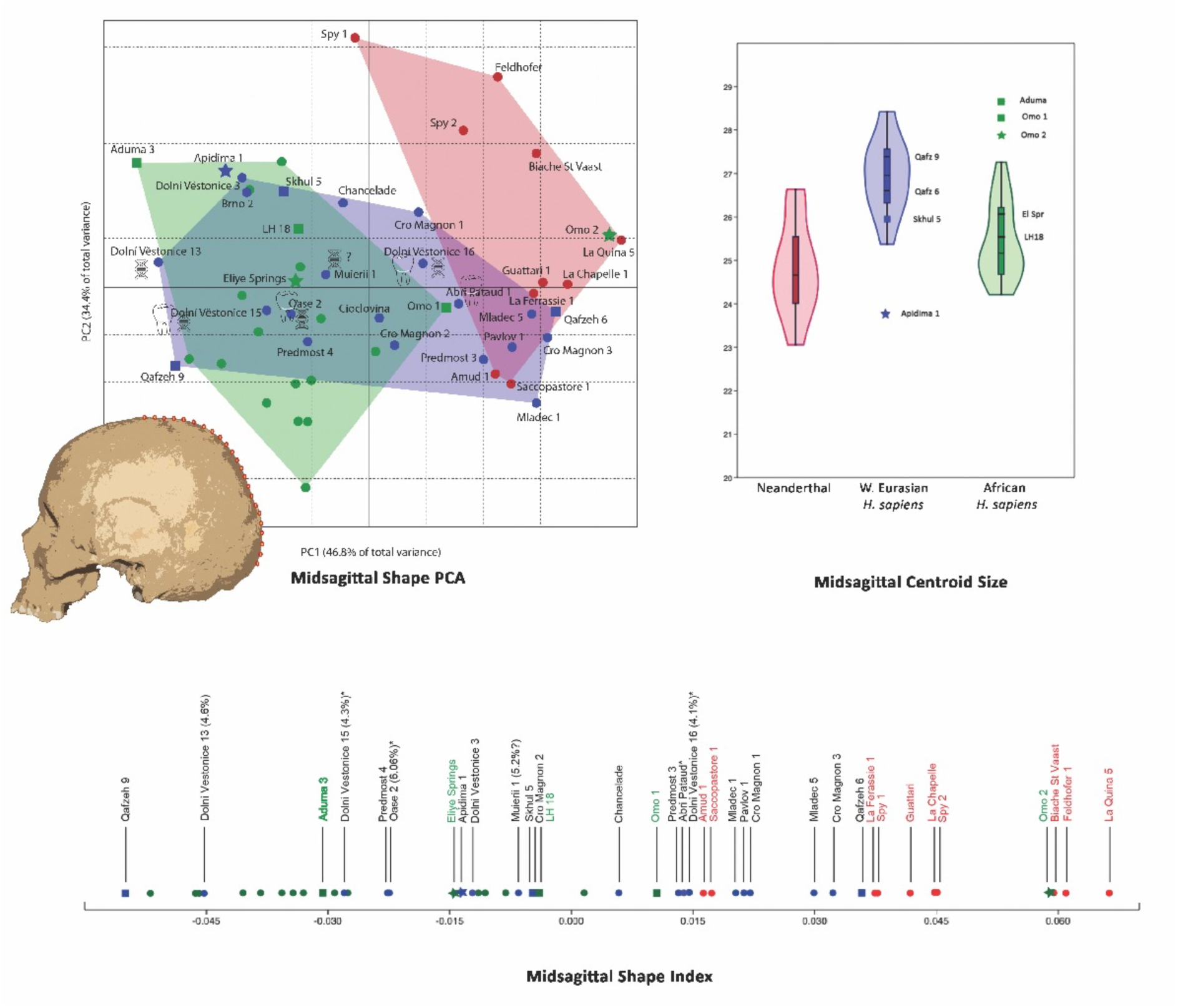
Posterior cranial (midsagittal) profile analysis. Top left: Principal Components Analysis, landmark / semilandmark dataset on modern human cranium model. Top right: Violin plot centroid size by group, minimum to maximum values, superimposed box plot showing median and 25-75 % quartiles. Bottom: Shape index. Colors and symbols as in Figure 3. Individuals with genetic evidence for hybridisation are marked with a DNA symbol in the PCA plot, with % Neanderthal ancestry given in the shape index. Individuals with atypical dental or sutural variation as reviewed in^132^ are marked with a TOOTH symbol in the PCA plot and with an (*) in the shape index.

In contrast to the African samples, multiple Eurasian *H. sapiens* specimens overlap with (Qafzeh 6, Pavlov 1, Mladeč 5, Cro Magnon 3), or plot close to (Cro Magnon 1, Mladeč 1, Abri Pataud 1, Predmost 3) the Neanderthal convex hull. Many also show Neanderthal-like shape indices. Some of these individuals have previously been described as possessing occipital ‘hemibuns’, posterior projections of the occipital bone reminiscent of those shown by Neanderthals, possibly due to Neanderthal ancestry. Unfortunately, no genomic evidence is available for them. Elongated cranial profiles in fossil *H. sapiens* might also result from the retention of ancestral morphology represented here by Omo 1 and possibly Omo 2. However, the Omo specimens greatly predate both the Levantine and the European Upper Paleolithic samples – by ca. 60-90 ky and >160 ky, respectively – making recent admixture a more likely explanation for the observed variation in the Upper Paleolithic, and perhaps also the Near Eastern sample, a possibility that requires further investigation. On the other hand, Oase 2, which exhibits upper third molar megadontia^83, 132^ and has also recently been found to have relatively elevated Neanderthal admixture (6.06 %;^124^), plots in the centre of the African convex hull in the PCA and shows a modern human-like shape index. So does Dolní Vestoniče 15, which shows 4.3 % Neanderthal ancestry^4^ as well as a conical mandibular supernumerary tooth in the region of the left canine root and rotation of the left mandibular premolar^132^; the supernumerary tooth in particular might be interpreted as possibly resulting from admixture (though being a more common form of supernumerary tooth it is not strong evidence).

Furthermore, several additional specimens with known, relatively low, Neanderthal genetic components (Figure 4) have shape index values within the range of African *H. sapiens*. Finally, the proposed early *H. sapiens* Apidima 1 specimen plots with African *H. sapiens* both in the PCA as well as in the shape index, but is characterised by a smaller centroid size, consistent with retention of ancestral morphology as well as with possible admixture.

Although the hemimandible and posterior cranial profile analyses are broadly consistent with expectations based on animal hybrid models, the facial dataset shows a different pattern (Figure 5). Here the African *H. sapiens* sample (with the exception of Late Pleistocene specimen Hofmeyr) falls within the more dispersed shape space of the Eurasian *H. sapiens* group; both plot away from the tightly clustering Neanderthal sample. The two *H. sapiens* samples also show roughly equivalent centroid sizes. The Eurasian sample, however, is considerably more variable in shape, as reflected in their more widely diverging PC scores, with most specimens, including all individuals with known Neanderthal genetic components, falling outside of the African *H. sapiens* convex hull (i.e. transgressive relative to African *H. sapiens*). Such increased variability is consistent with an admixed sample, but could also result from sampling bias, a greater temporal variability in our Eurasian *H. sapiens* sample, or from within-species geographic variation – though similar temporal and geographic variation did not lead to this pattern in the other analyses. Similarly, the facial shape index values of the African *H. sapiens* specimens fall within the Eurasian *H. sapiens* range and away from that of Neanderthals. Again, there is no relationship between the facial shape index and the percentage of Neanderthal ancestry in the specimens for which the latter is known (Figure 5). The early modern humans from the Near East, Qafzeh 9 and Qafzeh 6, plot in more intermediate positions in the PCA, and have intermediate facial shape indices, although still clearly away from the Neanderthal range.

**Figure 5:**
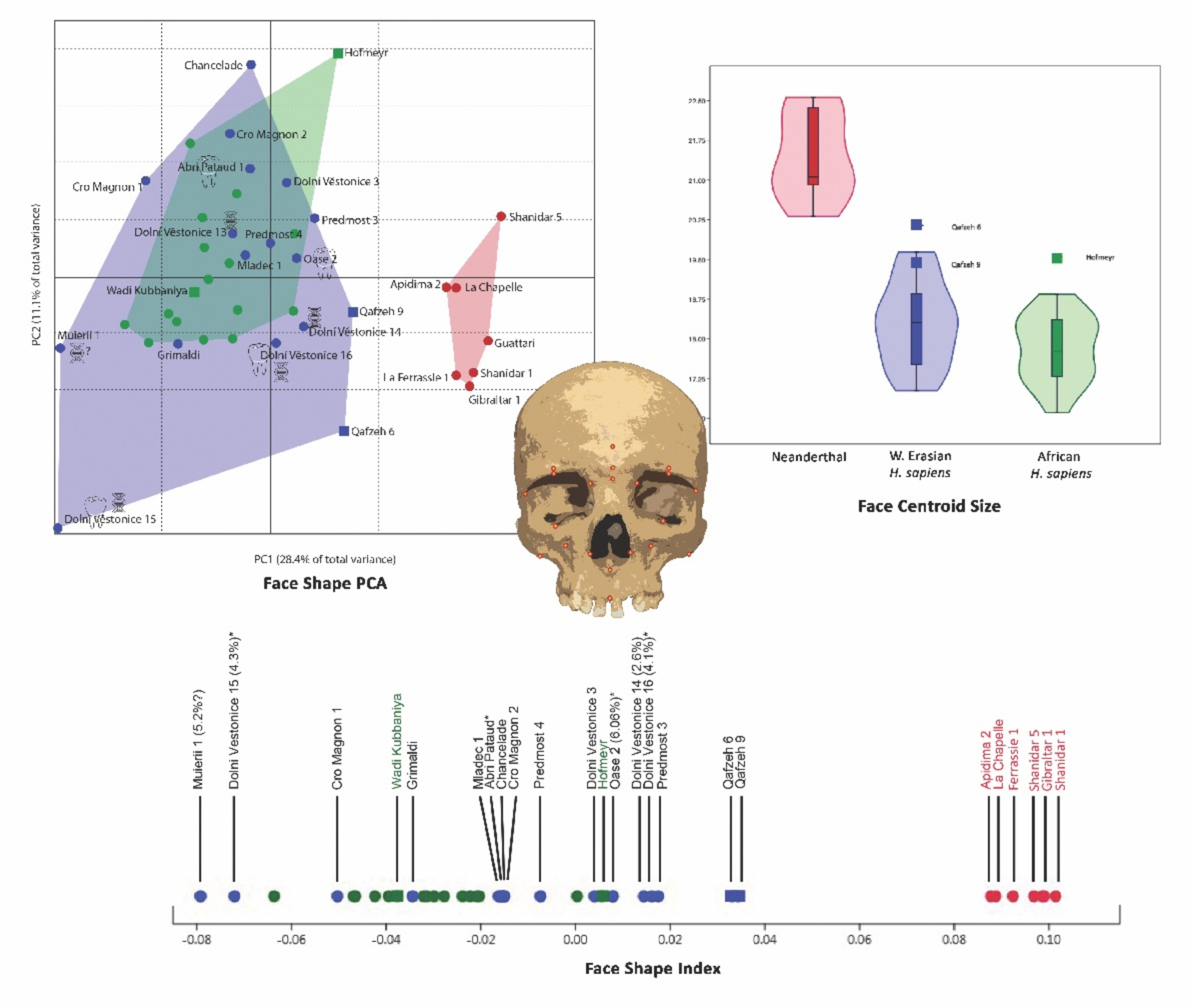
Face analysis. Top left: Principal Components Analysis, landmark dataset shown on modern human cranium model. Top right: Violin plot centroid size by group, minimum to maximum values, superimposed box plot showing median and 25-75 % quartiles. Bottom: Shape index. Colors and symbols as in Figure 3. Individuals with genetic evidence for hybridisation are marked with a DNA symbol in the PCA plot, with % Neanderthal ancestry given in the shape index. Individuals with atypical dental or sutural variation as reviewed in^132^ are marked with a TOOTH symbol in the PCA plot and with an (*) in the shape index.

## Discussion and Future Considerations

We did not approach this study by asking whether hybridization was common in Late Pleistocene Europe – though current evidence suggests that it may have been. Instead we wanted to evaluate how admixture manifests in the skeleton and whether different lines of evidence, morphological as well as genetic, can help reveal the presence of admixture, making it possible to identify hominin hybrids on the basis of either (or both). That being said, our evaluation of the morphology of Late Pleistocene Eurasian *H. sapiens* against predictions based on model organisms is based on small and imperfect samples (i.e. poor representation of African early *H. sapiens*; few individuals with both genomic and morphological data available; representation of primarily later rather than early generation hybrids). Recent African individuals are also imperfect models for early *H. sapiens* given that they have gone through their own process of evolution relative to the population for which we are using them as a proxy. Furthermore, the interpretation of the observed patterns is complicated by equifinality, as phenotypic variation consistent with admixture may also result from other processes, especially retention of ancestral features, and by sampling limitations which may underestimate the true variability of the groups included in our analyses. We therefore can only provide tentative and preliminary answers to the questions posed. These answers, nevertheless, can form the basis for future work exploring hybridization in the human fossil record.

To summarize, we explored whether our Late Pleistocene Eurasian *H. sapiens* sample fits our predictions for a Neanderthal-early *H. sapiens* population with a history of hybridisation. For our mandibular and posterior cranial datasets, we found that it was intermediate in shape and size between Neanderthals and African *H. sapiens* and transgressive in aspects of shape, patterns consistent with hybridization across the sample as a whole. Facial shape, on the other hand, did not provide a clear signal, although the large variation in the Eurasian *H. sapiens* sample and high proportion of transgressive individuals is also consistent with hybridisation in that dataset. It is unclear why different anatomical regions would differentially preserve the signal of hybridisation, if indeed that is the signal being detected here. Facial and mandibular shape has been argued to be affected differentially by selection and by adaptive or plastic responses to external, environmental factors (e.g.^128–130^). Facial morphology is also widely recognised as important in species recognition and social interactions among primates^131^, and may therefore be under selective pressure to conform more closely to the backcrossing population. Finally, in all of our analyses Late Pleistocene Eurasian *H. sapiens* as a sample were closer in shape to African *H. sapiens* than to Neanderthals, as expected under conditions of asymmetric gene flow (hypothesized for large differences in parental population sizes, as postulated for early modern Europeans relative to late Neanderthals, e.g.^120^), or, more importantly, for a sample comprising multiple, later generation (i.e. more backcrossed into modern humans) hybrids.

In terms of individual specimens, no direct relationship was found between estimated levels of Neanderthal ancestry based on genomic evidence, where known, and anatomical shape/size, nor between this genomic evidence and expression of developmentally abnormal dental or sutural features as reported in the literature. This was the case also in the early generation Neanderthal-modern human hybrid, the mandible Oase 1, whose only obvious phenotypic signals of hybridization are its very large overall size and megadontia. This result is perhaps not surprising, as estimated admixture percentages may vary across most specimens due to noise or sequencing depth.

Furthermore, the critical factor for the expression of Neanderthal-like features is most likely the presence of particular alleles relevant for the expression of specific phenotypes, rather than overall percentages of Neanderthal ancestry (as has recently been argued for cranial globularity by^35^).

Assuming that cranio-mandibular morphology is at least in part under genetic control, the comparatively moderately elevated Neanderthal genetic component shown by, e.g., Dolní Vestoniče 16, may comprise alleles influencing development of the masticatory region and neurocranium, which resulted in shape similarities to Neanderthals reflected by this specimen’s mandibular and midsagittal profile shape indices and PC scores (Figures 3, 4) and in the known misalignment of the maxillae along the intermaxillary suture^136^ but not in facial similarities (Figure 5).

Are these skeletal morphologies useful predictors of admixture in samples where no genetic evidence is available? At the moment the patterns observed when considering a larger sample/population are the most informative. As regards individual specimens, the signals are often mixed, even across anatomical regions for the same individuals, likely reflecting differential preservation of the hybridisation signal according to anatomical region (see above). The state of preservation and degree of completeness of a fossil, therefore, may influence whether an admixture signal can be detected. This signal will likely further be influenced by the differential expression of Neanderthal-like or developmentally abnormal features according to the presence of particular Neanderthal alleles or the degree and/or recency of ancestry. Nevertheless, some observations can be made. The individuals Cro Magnon 3, Pavlov 1, and Mladeč 5 are the only ones across all our analyses that plot well within the Neanderthal convex hull in the PCA and show Neanderthal-like shape index values in their posterior midsagittal cranial outline, the only dataset in which they could be included. On this basis we may hypothesize that they have a Neanderthal genetic component comprising alleles important for cranial shape. A similar argument could be made for the Qafzeh 6 individual, who, in contrast to Qafzeh 9 from the same site, also overlaps with Neanderthals in the posterior cranial analysis. The Qafzeh specimens are also the only ones that show a somewhat intermediate position in the facial analysis. These results, together with the high levels of variation in one site, and the geographic origin in the Levant, a postulated contact area between Neanderthals and modern humans^5^, raise the possibility that the Qafzeh individuals may also have some Neanderthal ancestry^84^. Even though such indications are intriguing, they cannot be considered conclusive and must be treated as hypotheses, especially since, as mentioned above, similar phenotypes might be consistent with different underlying causes. Nevertheless, it is possible to evaluate the likelihood of such alternative explanations on a case-by-case basis. For example, because a rounded cranium is a derived *H. sapiens* feature, an alternative hypothesis for a relatively elongated cranial phenotype could be that it results from retention of the ancestral, elongated condition. An ancestral retention, however, is more convincing for Qafzeh, which represents an early *H. sapiens* population dating to ca. 100-130 ka (Table 1), than for the Late Pleistocene European specimens, which greatly postdate the establishment of the derived condition^117, 137^.

Finally, recent suggestions that skeletal anomalies in some Upper Paleolithic and Neanderthal samples result from inbreeding^132, 138^ may further complicate the interpretation of developmental abnormalities as indicators of admixture. Indeed both processes are expected to have taken place in the highly dynamic conditions of cyclical environmental change of Pleistocene Eurasia, which likely resulted in repeated isolation of populations in refugia areas, sometimes leading to local extinctions, but also to population expansion and dispersals^139^. Under these conditions, paleodemes have been proposed to resemble ‘tidal islands’, often isolated but occasionally flooded with expanding / dispersing populations and their genetic material^139^.

However, although empirical evidence from primates for the skeletal expression of inbreeding is limited, the evidence that does exist suggests that it is associated with abnormalities (e.g. reduced size, anencephaly, polydactly, syndactly, limb malformations^140–144^) that are different from those shown to occur in hybrids (e.g. increased size, extremely rare dental and sutural traits with no other associated diseases or syndromes^1, 84, 96, 97^). This indicates that it should be possible to distinguish between these causal phenomena, and their relative contributions to the morphology we see in the fossil record, going forward.

This study compared genomic and morphological data sets, in order to interrogate the fossil evidence for Late Pleistocene hybridisation between Neanderthals and early *H. sapiens*, for which we currently have substantial evidence. We urge further studies of the phenotype to expand our ability to detect the ways in which migration, interaction and genetic exchange have shaped the human past, beyond what is currently visible with the lens of ancient DNA. It is particularly important to examine such data sets together to understand the effects of hybridization on the morphology of later generation hybrids, and whether these effects vary by anatomical region. The results provided here should form the basis for developing hypotheses to be tested against the human fossil record in the future.

## Acknowledgements

This research was supported by the European Research Council (ERC CoG no. 724703 CROSSROADS), the German Research Foundation (DFG FOR 2237 ‘Words, Bones, Genes, Tools), and the National Research Foundation of South Africa (Grant# 117670). We thank all curators and institutions that allowed us access to the fossil specimens used in our analyses, K. Warren for collecting and analysing the data used in Figure 1, A. M. Bosman and C. Röding for help with processing the datasets used in Figures 3-4, H. Rathmann and J. Beier for help with the figures, and C. Posth for important feedback. We are grateful to three anonymous reviewers whose comments and suggestions greatly improved our manuscript.

## TEXT BOX

### Are Neanderthals Distinct Species and Does It Matter?

There is a longstanding debate around the taxonomy of Neanderthals and especially whether they represent distinct species from *Homo sapiens*. Even the two authors of this manuscript differ in opinion on this matter. Applying species concepts that rely on complete reproductive isolation (e.g.^145^), where species are defined in part according to their ability to interbreed and produce fertile offspring, it is now clear that they would be considered the same species. However, such species concepts are of little value in the context of hybridising lineages. Instead, most evolutionary biologists would agree that species are defined by retained morphological, behavioural and genetic differences even in the face of gene flow^146^, a definition that more clearly argues for species- distinctiveness for Neanderthals (and possibly others; e.g.^67, 147^). However, as pointed out elsewhere^148^, this definition also implies an ability to coexist geographically without the fusion of lineages^149^, which does not appear to be the case between Neanderthals and *H. sapiens*. Although these lineages appear to have evolved their differences initially in geographic isolation, post-contact gene flow resulted in the subsequent disappearance, at least morphologically, of all but one lineage, *H. sapiens*. This may suggest that Neanderthals and *H. sapiens* may not have been able to coexist without lineage fusion^148^. However, the disappearance of Neanderthals and their relatively low genetic contribution to later humans may also point to competitive exclusion, in the context of reported reduced hybrid fitness (e.g.^122^), environmental instability and cultural / demographic differences (e.g.^150, 151^). Given these differences of opinion, whether these taxa represent genomically coherent, monophyletic species remains an open question, one which may be resolved in the future as more and more genomes are sequenced across the full distribution of the taxa. More importantly, the question of specific status is irrelevant for the investigation undertaken here. Regardless of whether Neanderthals, *H. sapiens* and others represent distinct species, they are clearly genetically differentiated lineages. How different lineages need to be for gene exchange to be categorised as hybridization is subjective. Typically, however, it is at or above the rank of subspecies^152^, which is almost certainly the case here. This is also the case for the extant taxa used as models for considering the morphological effects of admixture. Baboons are particularly interesting in this respect due to their extensively documented phenotypic, ecological and behavioral distinctiveness combined with their ability to interbreed extensively (see^40^). The history of disagreement over specific versus subspecific levels of distinctiveness among baboon ‘allotaxa’ broadly parallels the discussion surrounding Neanderthals and *H. sapiens*, pointing to the shortcomings of species concepts as applied not only to the fossil record, but also to extant mammalian taxa.

